# Humidity shapes the thermal niche of *Anopheles stephensi*, an invasive malaria vector

**DOI:** 10.64898/2026.03.28.715035

**Authors:** Britny Johnson, Paul J. Huxley, Joel J. Brown, Brandon D Hollingsworth, Eric R. Bump, Brandyce St. Laurent, Jared Skrotzki, Leah R. Johnson, Mercedes Pascual, Michael C. Wimberly, Ajeet Mohanty, Courtney C. Murdock

## Abstract

Vector-borne pathogens cause 17% of all human infectious diseases, and rising global temperatures are shifting the distribution and abundance of mosquito vectors. Because mosquitoes are ectotherms, temperature strongly governs biological rates and physiology; however, mosquitoes also experience other environmental factors that may interact with temperature to shape the thermal performance of traits driving population dynamics. Here, we use a factorial life-table experiment spanning five relative humidities (30–90%) and seven temperatures (16–38*^◦^*C) to show that humidity modifies the thermal performance of key fitness traits in adult *Anopheles stephensi*, an invasive urban malaria vector. When integrated into a demographic model, humidity markedly reshapes projections of population fitness relative to temperature-only models, suppressing growth and contracting year-round suitability in hot, arid regions while enhancing fitness in more humid or high-elevation climates characteristic of South Asia and Africa. Together, these results highlight the need to integrate multiple environmental drivers into projections of climatic suitability, as temperature-only approaches may mischaracterize both the magnitude and spatial structure of mosquito population fitness. More broadly, our findings demonstrate how moisture availability reshapes thermal niches, population fitness, and climate-driven projections of vector distributions.

## Introduction

In the broader field of ecology, significant attention has been devoted to understanding how temperature influences the distribution and abundance of organisms, driven by concerns over climate change and the impacts of increasing global temperatures (Walther et al., 2002). This is particularly important for vector ecology, where vectorborne pathogens account for 17% of all human infection illnesses, causing approximately 700,000 deaths annually (Venkatesan, 2024), while also placing significant constraints on global agriculture and wildlife health (Aguirre, 2017; Stuchin et al., 2016). As most disease-causing vectors are ectotherms, their fitness and survival are directly influenced by variation in ambient temperature (Mordecai et al., 2019), and numerous studies have demonstrated that variation in ambient temperature is one of the most fundamental environmental factors constraining the geographical distribution and seasonal dynamics of arthropods like mosquitoes (Paaijmans et al., 2013; Villena et al., 2022). However, ectotherms experience a complex suite of other environmental factors, both abiotic and biotic, that vary their effects on fitness and likely interact to shape their current and future distribution and abundance (Huxley et al., 2022; Kleynhans and Terblanche, 2011; Liu and Gaines, 2022). Therefore, to better predict shifts in mosquito ranges, survival, and invasion potential in a rapidly changing environment, it is essential to consider how multiple environmental factors affect mosquito fitness.

The relationship between mosquito life history traits and temperature is often characterized by laboratory experiments that place mosquitoes across a range of temperatures and measure the effects of temperature on mosquito life history traits that govern mosquito population and disease dynamics. Temperature-trait relationships follow a nonlinear, unimodal Thermal Performance Curve (TPC) that is bound by a minimum temperature (*T*_min_) and maximum temperature (*T*_max_) that represent the operative range of a given physiological process, and an intermediate temperature (*T*_pk_) where trait performance is maximized (Angilletta Jr, 2009; Huey and Kingsolver, 1989). Thermal performance curves are consistent with the metabolic theory of ecology, where metabolic efficiency is predicted to increase as temperatures increase but then will decline after exceeding the *T*_pk_ due to compromised protein stability, cell membrane integrity, and neuromuscular coordination, with organismal death occurring at the *T*_max_ (Kingsolver and Woods, 2016). Thermal performance curves have been widely used to predict current and future environmental suitability for a diversity mosquito species and the pathogens they transmit (Villena et al., 2022; Ryan et al., 2023; Miazgowicz et al., 2020; Mordecai et al., 2019, 2013), as well as other ectothermic organisms.

In addition to temperature, water availability is a critical yet often overlooked environmental variable constraining the distribution and abundance of ectotherms, like mosquitoes (reviewed in Brown et al. (2023)). While temperature sets the rate of metabolic reactions, water is the universal solvent in which all cellular processes occure (e.g., nutrient transport, osmoregulation of electrolytes, etc.) (Brown et al., 2023; Chaplin, 2006; Cheuvront and Kenefick, 2014; Chown, 2002; Rozen-Rechels et al., 2019). Thus, the optimal regulation of both body temperature and water balance is crucial for organismal fitness (Bradshaw, 2003). Due to the fundamental relationship that exists between temperature and the amount of moisture the air can hold, variation in both relative humidity and temperature will alter the degree of moisture stress ectothermic organisms, like mosquitoes, experience. For a given amount of atmospheric moisture, warmer temperatures will increase the total amount of water the air can hold, increasing the vapor pressure deficit and the amount of potential water loss an organism experiences. As temperatures warm, variation in relative humidity can either buffer (high relative humidity) or exacerbate (low relative humidity) the negative effects of high temperature on mosquito fitness Brown et al. (2023). The current way in which TPCs are characterized does not account for these effects. Even when relative humidity is held constant, increases in temperature will increase the vapor pressure deficit and evaporative stress organisms experience. Thus, it is currently unclear if the CT_max_ of a given trait is driven by the effects of temperature on metabolic function, or rather, is a function of dehydration and water stress. Understanding this interaction and the physiological mechanisms underpinning mosquito responses to abiotic constraints is critical for predicting mosquito distributions and population dynamics now, and in the future in response to global change (Deutsch et al., 2008; Pörtner and Farrell, 2008; Dillon et al., 2010).

*Anopheles stephensi* has historically driven large malaria outbreaks in cities of India, Iran, and Pakistan and has now invaded seven countries in Africa (Mnzava et al., 2022; Sinka et al., 2020; Taylor et al., 2024) threatening ongoing malaria elimination efforts. Following its invasion in Djibouti, malaria cases have surged more than 2000-fold. This study builds on our prior work showing that relative humidity reshapes the thermal performance of juvenile traits in *An. stephensi* (Huxley et al., 2025). (Venkatesan, 2024; Vogels et al., 2023). Here, we extend that stage-specific framework to the adult life stage by quantifying how temperature (16°C–38°C) and relative humidity (30%–90%) variation jointly affect adult lifespan, fecundity, and biting rate. We then integrate these adult trait responses with previously estimated juvenile trait relationships to calculate humidity-dependent maximal population growth rate (*r*_m_), thereby linking humidity effects across life stages within a single demographic framework. We hypothesize that decreases in relative humidity as temperatures warm will increase the amount of desiccation stress a mosquito experiences, resulting in a reduction in the predicted *T*_max_ that limits trait performance and the *T*_pk_ where performance is maximized. We find relative humidity variation to significantly affect the thermal performance of *An. stephensi*, as well as our predictions for overall environmental suitability.

## Methods

### Experimental Design

Urban-type form *Anopheles stephensi* mosquitoes acquired from a long-standing colony (∼40 years) at Walter Reed Army Institute of Research, via the University of Georgia, were maintained under standard rearing conditions using established methods described in Miazgowicz et al. (2020). For the experiment, mated females (3-to-5 days old) were provided a whole human bloodmeal (O+ male donor 25–65 yr, Bio IVT) for approximately 30 minutes via a water-jacketed membrane feeder. Individual blood-fed females were then randomly distributed into separate containers (16 oz. paper cup; mesh top) and assigned to one of seven constant temperatures (16°C, 20°C, 24°C, 28°C, 32°C, 35°C, 38°C ± 0.5°C) across five relative humidities (30%, 45%, 60%, 75%, 90% ± 0.5%) in a full factorial design combined with a small initial pilot run across a subset of conditions for a total of 44 temperature-humidity combinations and 20 females per treatment (*n* = 700 individual females). Following mating and blood-feeding, females were housed individually and monitored separately.

Each container was provided with an oviposition site (plastic 2 oz. cup and Whatman filter paper) moistened with reverse osmosis water. Individual females were offered a bloodmeal for 15 min each day and were scored as bloodfed through visual verification of the abdomen immediately after feeding. Oviposition sites were rehydrated and checked for eggs daily, and females were followed until all mosquitoes died. Across treatments, all individuals had died by day 60 with the exception of three separate instances where a single individual was still alive. Therefore, in our analysis, we considered all individuals to have died by the end of the experiment. From these data, we quantified the effects of variation in relative humidity on the thermal performance of adult lifespan, fecundity, and biting rate. The temperature-trait relationships for these adult traits were then used to update a model for the maximal population growth rate (*r*_m_) (Cator et al., 2020; Huxley et al., 2025), to assess how variation in relative humidity impacts the thermal performance of this key metric of population fitness.

Variation in temperature (±0.5°C) and relative humidity (± 5% RH) were tightly regulated across Percival incubators. Thus, because incubator-level differences were negligible relative to treatment effects, individually monitored females were treated as biological replicates. To control for any possible microclimatic variation within an incubator, individually housed mosquitoes were randomly rotated across shelves and positions on shelves. Each day, blood-feeding order for each treatment was randomized to avoid feeding order bias. Finally, to achieve an experiment of this size, treatments were blocked, with environmental treatments randomly assigned across different cohorts of mosquitoes.

### Fitting temperature-trait relationships in the context of relative humidity

We fit thermal performance curves (TPCs) for the adult life history traits measured across relative humidity levels using a Bayesian inference framework (Shocket et al., 2025; Huxley et al., 2025; Johnson et al., 2015). Adult lifespan was modeled using a temperature-dependent median function with a Weibull likelihood, implemented in JAGS as the *a* = 1 special case of the generalized gamma family (Supplementary Eqns. 2–4). This formulation yields flexible, unimodal, and potentially asymmetric thermal performance curves. In this formulation, the median lifespan is parameterized by the temperature of peak performance (*T*_pk_), the peak trait value (*B*_pk_), a curvature parameter controlling the rate of decline away from *T*_pk_, and a shape parameter governing asymmetry in the rise and fall of the curve. We selected this approach because it permits long tails at the extremes of the temperature range, which is consistent with the biology of adult survival: absolute lower and upper thermal limits cannot be confidently estimated due to the absence of data at cold temperatures (below 10°C) and the persistence of survival even at the highest experimental temperatures. We chose this temperature-dependent median model with Weibull likelihood over a quadratic alternative (often used in modeling mosquito lifespans; e.g., Mordecai et al., 2019) because it consistently outperformed the quadratic model in comparisons using the Watanabe–Akaike Information Criterion (WAIC; Supplementary Fig. 1; Watanabe, 2010; Gelman et al., 2014). The model was implemented in R (R Core Team, 2025) using the `rjags` package (Plummer, 2025), with posterior inference obtained from Markov chain Monte Carlo (MCMC) sampling. Estimates of *T*_pk_ and *B*_pk_ were obtained numerically from the posterior distributions for each humidity level’s fitted curve and summarized using posterior medians and Highest Posterior Density (HPD) intervals.

We modeled the fecundity rate and biting rate thermal performance curves (TPCs) for each humidity level by fitting the standard Brière model (Supplementary Equation 1; implemented in the `bayesTPC` package in R; Briere et al., 1999; Sorek et al., 2025). Daily fecundity (*b*) was calculated for each female as the total number of eggs laid divided by the number of days the female was alive (eggs × female*^−^*^1^ × day*^−^*^1^). To characterize the thermal dependence of maximal egg-production capacity among ovipositing females, we fit Brière TPCs using females with positive fecundity rates only (*b >* 0) and extracted *b*_max_ and *T*_pk_ numerically from posterior draws. Consequently, *b*_max_ should be interpreted as peak fecundity conditional on oviposition (i.e., among females that laid at least one egg), rather than the unconditional expected egg production across all females.

Biting rate (propensity to blood feed within 15-minute daily offering) was calculated as the lifetime number of blood meals for each individual divided by its adult lifespan. For both fecundity and biting rate, the temperatures at which performance peaks and its value at its peak (*T*_pk_ and *B*_pk_, respectively; Table 1) were estimated numerically from the posterior distributions for each humidity level’s TPC and summarized by the posterior medians and Highest Posterior Density (HPD) intervals.

**Table 1:** Definitions of model parameters.

| Parameter | Units | Description |
| --- | --- | --- |
| $r_m$ | day$^{-1}$ | Maximal population growth rate |
| $\alpha$ | days | Hatching-to-adult development time |
| $b_{\max}$ | eggs × female$^{-1}$ day$^{-1}$ | Peak per-capita daily egg production rate |
| $\kappa$ | day$^{-1}$ | Fecundity loss rate |
| $z$ | day$^{-1}$ | Adult mortality rate |
| $p_{\text{EA}}$ | probability | Survival averaged across juvenile stages |
| $T_{\text{pk}}$ | $^{\circ}\text{C}$ or K | Temperature at which trait performance peaks |
| $T_{\text{min}}$ | $^{\circ}\text{C}$ or K | Lower temperature at which trait performance ceases |
| $T_{\text{max}}$ | $^{\circ}\text{C}$ or K | Upper temperature at which trait performance ceases |
| $B_{\text{pk}}$ | Measurement unit of trait | Trait performance achieved at $T_{\text{pk}}$ |
| $T_{\text{opt}}$ | $^{\circ}\text{C}$ | Temperature at which $r_m$ peaks |
| $r_{\text{opt}}$ | day$^{-1}$ | $r_m$ achieved at $T_{\text{opt}}$ |

### Calculating maximal population growth rate, *r*_m_

To examine how variation in relative humidity affects maximal mosquito population growth rate (*r*_m_) across the full life cycle, we combined the adult trait thermal performance curves estimated in this study with previously published juvenile trait thermal performance curves for development and survival from Huxley et al. (2025). Specifically, we substituted the adult TPCs for lifespan and fecundity, *z* (1/lifespan) and *b*_max_, together with the juvenile TPCs for development time and egg-to-adult survival, *α* and *p*_EA_, into a continuous-time, stage-structured model based on the Euler–Lotka equation (Charnov, 1993; Savage et al., 2004; Amarasekare and Savage, 2012). Thus, the *r*_m_ calculations integrate humidity-dependent trait responses from both juvenile and adult life stages within a single demographic framework. Cator et al. (2020) derived an approximation suitable for the range of growth rates typically observed in arthropods (Eqn. 1; Table 1).

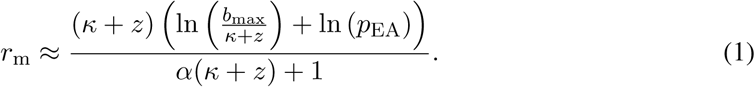

Here, *α* is the egg-to-adult development time (days), *b*_max_ is the peak reproductive rate (eggs × female^-1^ × day^-1^), *κ* is the fecundity loss schedule (individual^-1^ day^-1^), and *z* is the adult mortality rate (individual^-1^ day^-1^). Following Huxley et al. (2025), the term ln(*p*_EA_) replaces the original –*αz*_J_ term in Cator et al. (2020), allowing juvenile daily mortality to be approximated using the probability of emergence as an adult (egg-to-adult survival probability). As *r*_m_ has low sensativity to *κ* (Cator et al., 2020), we assume that *b*_max_ declines with age at a constant rate of 0.01 individual^-1^ day^-1^. Although Eqn. 1 is an approximation, it is sufficiently accurate provided *r*_m_ *<* 1 (in units of day^-1^; Cator et al., 2020), which generally holds for insect growth rates (Frazier et al., 2006; Pawar et al., 2024). Because it explicitly incorporates the underlying traits, Eqn. 1 can be used to analytically explore how variation in these traits influences *r*_m_.

Finally, to obtain all quantities of interest for *r*_m_ (*r*_opt_, *T*_opt_, *T*_min_ and *T*_max_), we used a similar procedure to the one that we used for adult fecundity rate. However, for *r*_m_ *T*_min_ and *T*_max_, we used the TPC posteriors at each humidity level to estimate the temperatures at which *r*_m_ was zero. Note that we refer to the temperature of peak *r*_m_ as *T*_opt_ (Table 1) rather than *T*_pk_ as we do for traits because the *T*_pk_ values for of those traits do not necessarily correspond to the thermal optimum of population fitness (optimal thermal fitness; Pawar et al., 2024).

### Trait sensitivity analysis

To determine the extent to which variation in humidity can affect the *relative* contributions of the adult fitness traits to *r*_m_’s temperature dependence we conducted a sensitivity analysis. This analysis uses the derivatives of *r*_m_ with respect to the traits of interest to determine the rate at which *r*_m_ changes with temperature (similarly to Mordecai et al., 2013; Cator et al., 2020). Full details of the approach are described in the Supplementary Information. Briefly, we derive an equation via the chain rule such that each summed term in the equation (Eqn. SE5) quantifies the relative contribution of each temperature dependent trait (described by the fitted TPC) in Eqn. 1 to the temperature dependence of *r*_m_ (i.e., to the overall derivative with respect to temperature). For simplicity and clarity in this calculation, we set the parameters in each TPC to the Maximum *A Posteriori* (MAP) estimator for each trait–humidity combination (Fig. 3) instead of using all posterior samples. The sample–based MAP estimator is calculated as part of the MCMC fitting process in the `bayesTPC` package (Sorek et al., 2025) in R.

### Mapping environmental suitability for mosquito population growth

To assess how temperature and relative humidity jointly shape the potential climatic niche of *An. stephensi*, we utilized high-resolution climate projections from the NASA Earth Exchange NEX-GDDP-CMIP6 archive. This global dataset contains bias-corrected simulations from multiple CMIP6 general circulation models at a 0.25°(∼25km) resolution. We extracted historical (1970-2000) runs for daily maximum and minimum temperature and specific humidity for all 23 available general circulation models. Daily maximum and minimum values were averaged to monthly means, and monthly mean temperature was computed as the average of the maximum and minimum temperatures. Monthly mean specific humidity was calculated and converted to relative humidity using the Magnus-Tetens equation, which incorporates saturation vapor pressure as a function of temperature. For each grid cell, the intrinsic population growth rate, *r*_m_, was computed using traitbased models. Two formulations were examined: a temperature-only model, which fixed relative humidity at 75% and a temperature × relative humidity model, in which both juvenile and adult trait responses varied with local relative humidity. Monthly *r*_m_ estimates were then aggregated into seasonal averages (January - March, April - June, July - September, October - December) to align with the north-south migration of the Intertropical Convergence Zone in Africa and the monsoon cycle in South Asia. We produced seasonal maps of *r*_m_ for Africa and South Asia under both models and calculated difference maps to identify where humidity altered population growth potential. Finally, we estimated the total area of year-round climate suitability by counting, for each pixel, the number of months with *r*_m_ > 0 and summing the area of pixels exceeding this threshold in all twelve months. These maps are heuristic tools to explore the sensitivity of *An. stephensi* climate suitability to humidity under realistic conditions within the species’ current geographic range and areas of potential future invasion. They are not meant to predict *An. stephensi* distribution or abundances.

## Results

### Relative humidity effects on temperature–dependent adult traits

The effect of temperature on adult trait responses was systematically shaped by variation in relative humidity, however these effects varied depending on the trait. Variation in relative humidity influenced the thermal performance of adult traits, with the direction and magnitude of this effect varying depending on the trait.

Generally, relative humidity did not alter the overall shape of the curves, with TPCs remaining qualitatively similar for lifespan (asymmetric, generalized gamma function, Fig. 1a), fecundity (asymmetric, Brière function, Fig. 1c), and biting rate (asymmetric, Brière function, Fig. 1f) across the range of relative humidity explored. For adult lifespan, we were unable to predict the effects of relative humidity on *T*_min_ or *T*_max_ with the generalized gamma function fitting the data best. However, qualitatively, increases in relative humidity appear to increase mosquito lifespan at cool and warm temperatures (Fig. 1a). In contrast, higher relative humidity decreased lifespan at temperatures that optimized this trait (*T*_pk_) with lifespan being approximately 16% higher at low relative humidity (45% RH) than high relative humidity (90% RH) (Fig. 1a, Table S1). However, credible intervals overlapped substantially for this TPC parameter, indicating limited evidence for differences among humidity levels (Fig. 1b; Table S1).

**Figure 1:**
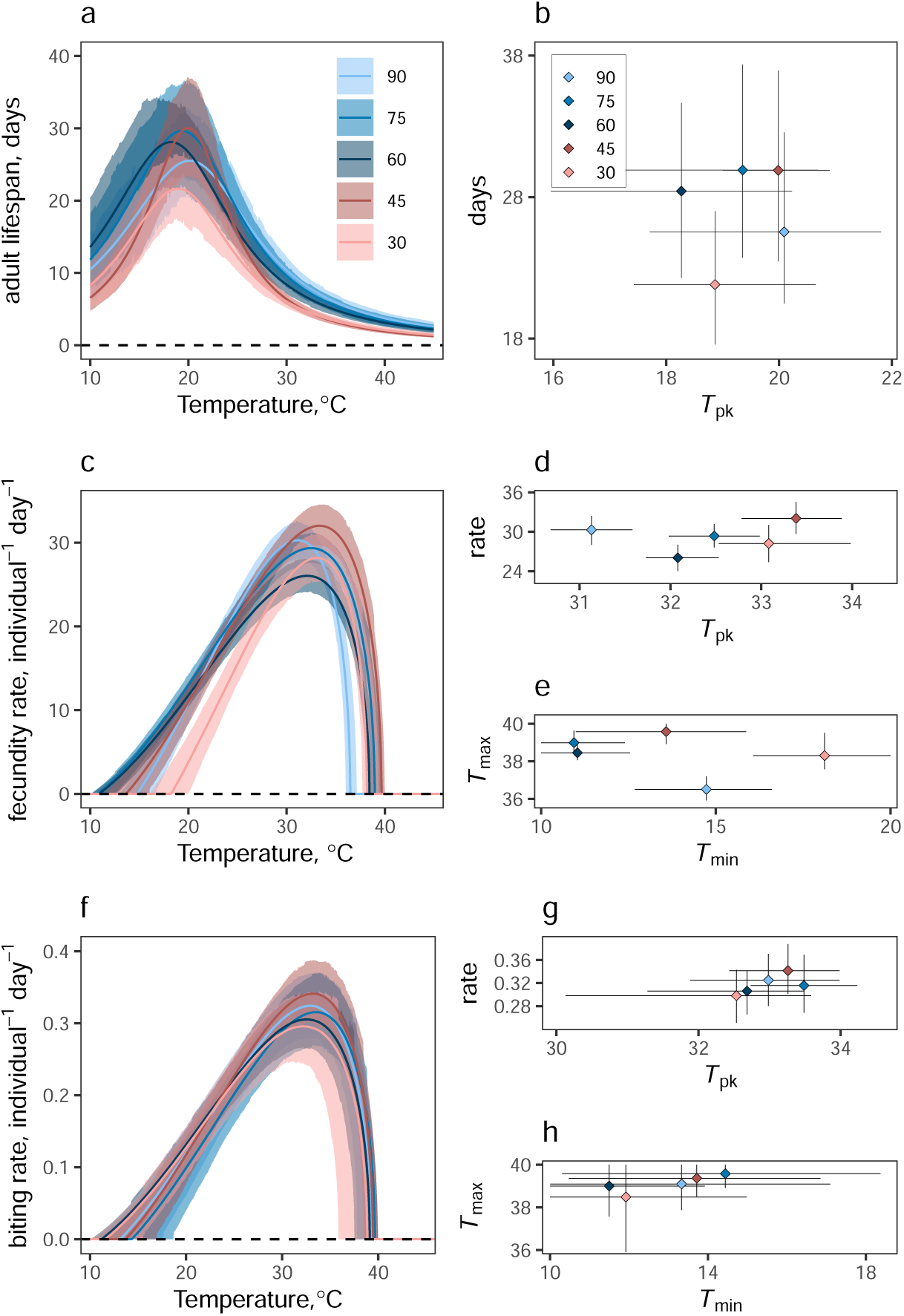
Relative humidity shapes the temperature dependence of adult fitness traits in *Anopheles stephensi*. The effects of relative humidity variation on the thermal performance curves (TPCs) of adult lifespan (1/*z* in Eqn. 1), fecundity rate (*b*_max_ in Eqn. 1), and biting rate (propensity to bloodfeed). Ball and stick diagrams represent how the predicted thermal peaks (*T*_pk_), and minimum (*T*_min_) and maximum (*T*_max_) vary at each relative humidity level for each trait. **Adult lifespan**, 1/*z*: **(a)** posterior distribution of fitted TPCs across humidity levels; **(b)** maximum lifespan vs. *T*_pk_. The adult lifespan TPCs were inverted to get *z* for the *r*_m_ calculations, which are shown in Fig. S2. **Fecundity rate**, *b*_max_: **(c)** posterior distribution of fitted TPCs across humidity levels; **(d)** maximum fecundity rate vs. *T*_pk_; **(e)** *T*_min_ vs. *T*_max_. **Biting rate**: **(f)** posterior distribution of fitted TPCs across humidity levels; **(g)** maximum biting rate vs. *T*_pk_; **(h)** *T*_min_ vs. *T*_max_. Shaded regions and error bars represent 95% HPD intervals summarizing posterior uncertainty.

For daily egg production (Fig. 1c), decreases in relative humidity generally shifted thermal performance toward warmer temperatures, with shifts in *T*_min_ and *T*_max_ by approximately 3 *^◦^*C (Fig. 1e, Table S2) and in *T*_pk_ by approximately 2 *^◦^*C (Fig. 1e, Table S2) as relative humidity decreased from 90% to lower humidity (30–45% RH) (Fig. 1c–e; Tables S2–S3). We also observed females laying the most eggs per day at extreme relative humidity (45% RH, ∼32 eggs day^-1^; 90% RH, ∼30 eggs day^-1^) and the fewest eggs at intermediate relative humidity (60% RH, ∼26 eggs day^-1^) (Fig. 1d, Table S3). Finally, the thermal performance of daily biting rate (Fig. 1f, Tables S4–S5) was largely unaffected by variation in relative humidity due to high variability across individuals in a female’s propensity to feed on a given day, which resulted in substantial posterior uncertainty across humidity levels (Fig. 1g–h).

### Relative humidity effects on temperature–dependent mosquito population growth rates

Maximal population growth rate (*r*_m_) showed a unimodal, asymmetric relationship with temperature across all humidity levels and remained positive above ∼14.5 *^◦^*C and below ∼40 *^◦^*C, with optima between 0.36 and 0.41 (Fig. 2a,b; Table S6–S7). Similar to the adult life history traits, variation in relative humidity systematically shifted the thermal performance of (*r*_m_) via humidity-driven changes in the underlying trait performance curves (Fig. 2a; Tables S6–S7). As relative humidity decreased from 90% RH to 30% RH, the predicted *T*_min_ increased by ∼3.5 *^◦^*C (Fig. 2c; Table S7), *T*_opt_ increased by ∼2.4 *^◦^*C (Fig. 2b; Table S7), and *T*_max_ increased by ∼1.7 *^◦^*C (Fig. 2c; Table S7). At temperatures that maximized *r*_m_, growth rates were highest at intermediate relative humidity (45%, 60%, and 75% RH; Table S7). Most notably, at optimal temperatures, the highest population growth rates were observed at low relative humidity (45% RH, *r*_m_ = 0.4068) and the lowest at high relative humidity (90% RH, *r*_m_ = 0.3597), consistent with the adult TPC parameters over the same range.

**Figure 2:**
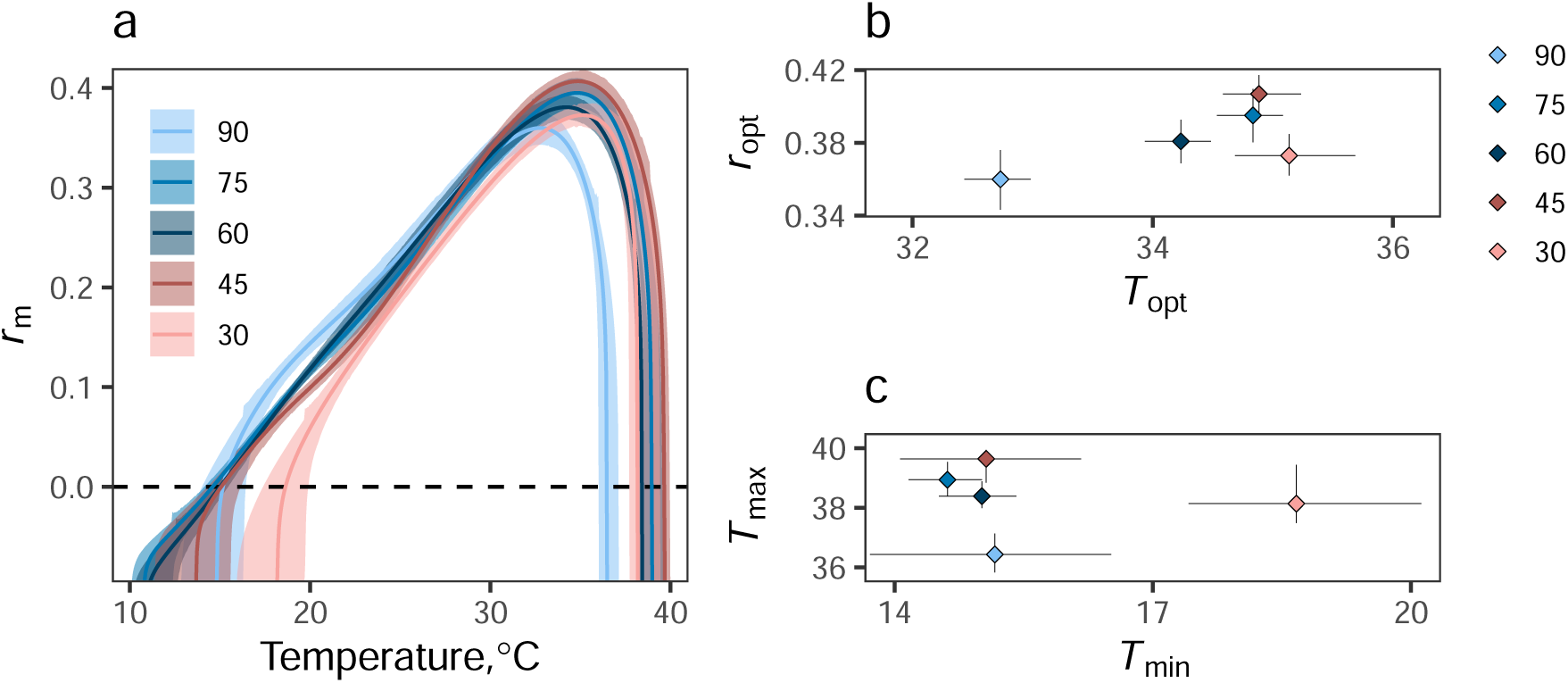
Effects of relative humidity on mosquito fitness traits shape the temperature dependence of maximal population growth rate, *r*_m_. (a–c) a. *r*_m_ TPCs across relative humidity levels. (b–c) b. *r*_opt_s versus *T*_opt_s, c. *T*_min_ versus *T*_max_ across humidity levels. Prediction bounds in **a** are HPD intervals calculated using the posteriors for each humidity–dependent TPC. Points (medians) in **b** and **c** were estimated numerically from the posterior distributions for each humidity level; bidirectional error bars show 95% Highest Posterior Density (HPD) intervals summarising posterior uncertainty.

### Sensitivity analysis

Variation in relative humidity had a non-linear impact on the sensitivity of *r*_m_, defined here as the rate at which *r*_m_ changes with temperature at a given temperature, driven primarily by the contribution of adult fecundity (*b*_max_) and, to a lesser extent, adult mortality rate (*z*), larval development rate (*α*), and pupae to adult survival (*p*_EA_) (Fig. 3). *r*_m_ is relatively insensitive to small changes in temperature at intermediate temperatures. However, sensitivity grows exponentially towards the extremes. Relative humidity modulates the range at which *r*_m_ is relatively insensitive to temperature, particularly at the low end, from approximately 20–38*^◦^*C at 30% RH to 12–39*^◦^*C at 60% RH to 18–36*^◦^*C at 90% RH. Adult fecundity is the primary driver of the rapid increase in sensitivity at extreme temperatures, due to its high sensitivity in these ranges. The contribution of adult lifespan is also high at the extremes, but its impact is small compared to fecundity. However, there is an increase in sensitivity of *r*_m_ to temperature from approximately 20–35*^◦^*C, driven by the contribution of adult lifespan.

**Figure 3:**
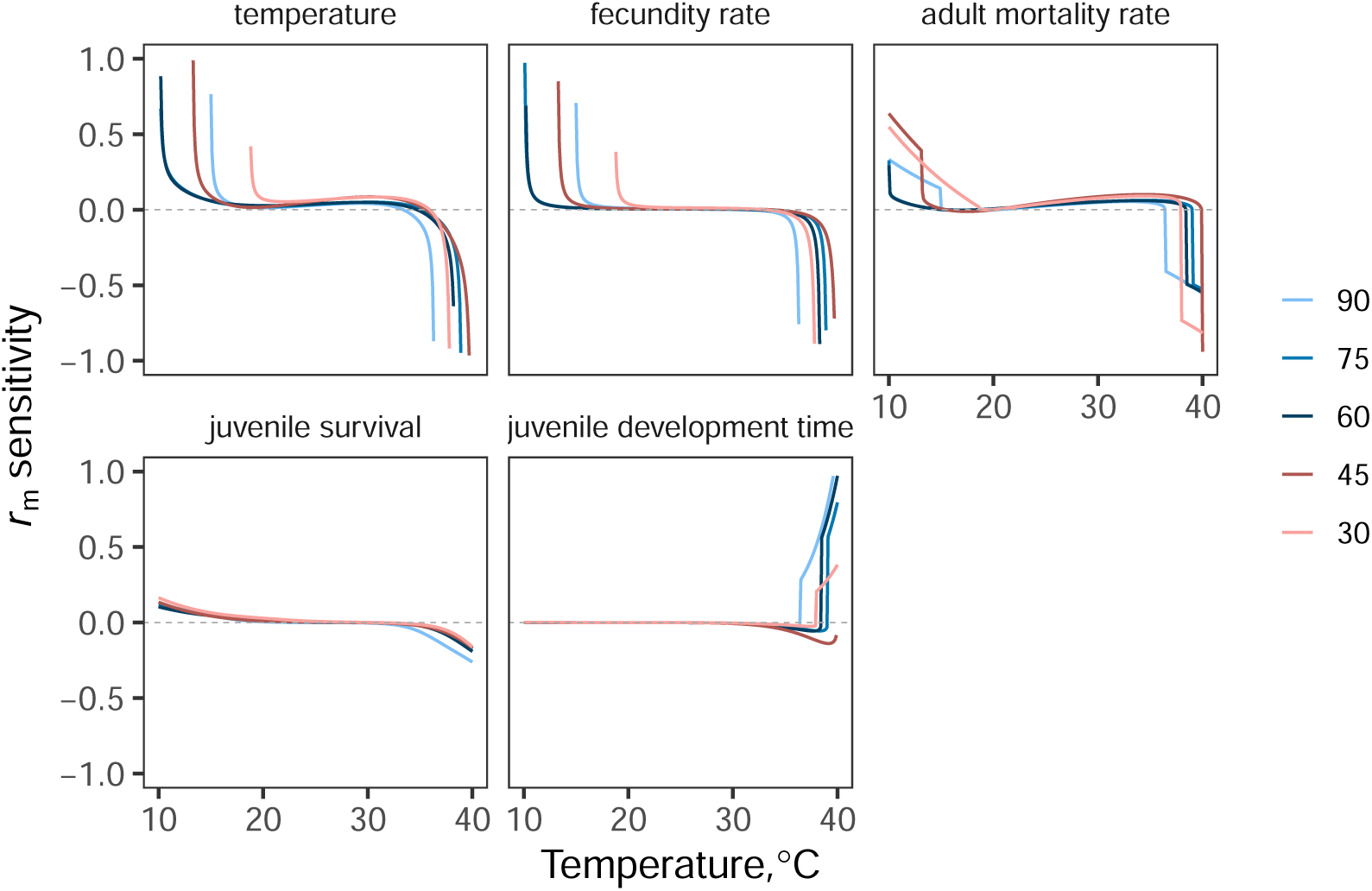
Sensitivity of maximal population growth rate, *r*_m_ to responses of underlying traits. Estimates of *r*_m_ are highly sensitive to temperature at extremes, driven primarily by the sensitivity of the maximum adult fecundity rate, (*b*_max_ in Eqn. 1, Table 1). This sensitivity is highest at the low and high temperature extremes, with relatively low sensitivity at intermediate temperatures. Relative humidity has a nonlinear effect on the sensitivity of *r*_m_ to temperature, particularly as relates to the low temperature where sensitivity increases exponentially. This occurs at approximately 19*^◦^*C for 30% humidity, 12*^◦^*C at 60 and 75% humidity, and 16*^◦^*C for 90% humidity. Adult lifespan and juvenile development time contribute most to the sensitivity of *r*_m_ to temperature at the extremes, <15*^◦^*C and >38*^◦^*C. However, the contribution of fecundity grows exponentially before reaching these temperatures, resulting in a relatively reduced impact on the sensitivity of *r*_m_ to temperature.

### Climatic suitability for mosquito population growth

Across both continents, the temperature-only model projected the most favorable growth conditions for *An. stephensi* (e.g., higher *r*_m_) in hot, lowland tropical climates, while much cooler higher elevation climates like the Himalayas, the Atlas Mountains, and the East African Highlands showed lower suitability (Fig. 4). In South Asia, including humidity effects led to higher *r*_m_ values across large swathes of the subcontinent, especially in central and southern India, Bangladesh, and Sri Lanka. At the same time, parts of Pakistan and the Himalayan foothills saw declines in *r*_m_ (Fig. 4). In Africa, the influence of humidity was more spatially varied. The inclusion of humidity effects suppressed *r*_m_ across much of the Sahara, Sahel, parts of the Horn of Africa, with localized increases in mountainous regions such as the East African Highlands (Fig. 4). These spatial patterns were seasonally dynamic. Humidity generally increased *r*_m_ during rainy seasons but reduced it during drier periods such as the South Asian pre-monsoon and African dry seasons. In terms of year-round climatic suitability, incorporating humidity reduced the size of the potential range. In South Asia, the year-round suitable area decreased from ∼3 million km^2^ (temperature-only) to ∼2 million km^2^, a reduction of ∼1 million km^2^. In Africa, it decreased from ∼24 million km^2^ (temperature-only) to ∼20 million km^2^ (temperature-relative humidity); a reduction of ∼4 million km^2^.

**Figure 4:**
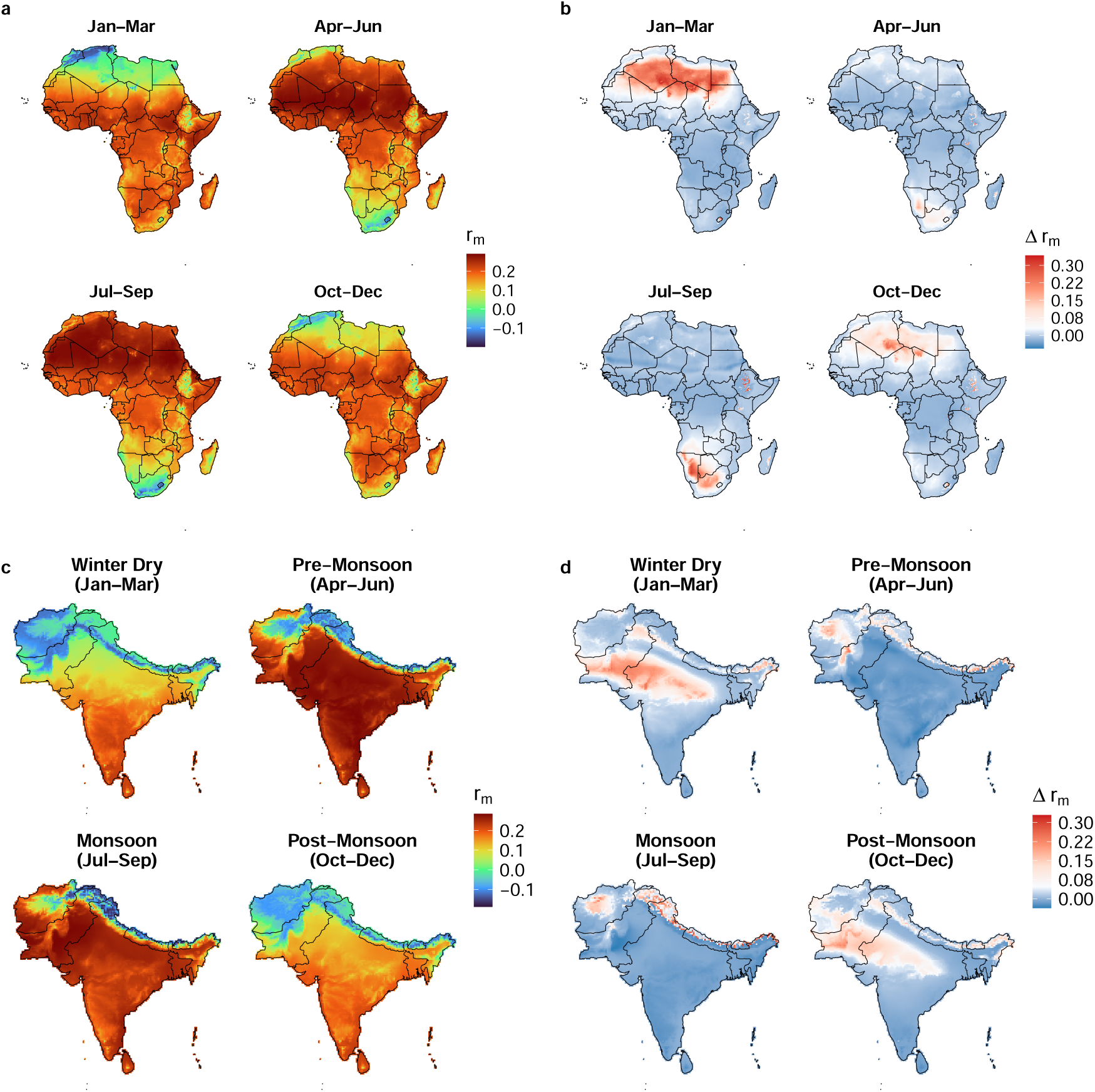
Seasonal spatial patterns of *r*_m_ derived from the NASA NEX-GDDP-CMIP6 daily historical dataset (1970–2000), for Africa (a and b) and South Asia (c and d) based on temperature and humidity. Maps (a) and (c) show the seasonal differences in *r*_m_ for the temperature-only model. In maps (b) and (d), Δ*r*_m_ = temperature-only *r*_m_ minus temperature- and humidity-dependent *r*_m_. Red (positive) values indicate that the temperature-only *r*_m_ model overestimated *r*_m_ and blue (negative) values indicate that temperature-only *r*_m_ model underestimated *r*_m_ (i.e., the temperature and humidity model predicted higher *r*_m_ than the temperature-only model). White values denote that both models were in agreement (Δ*r*_m_ = 0). Note, that these maps are intended to provide heuristic illustrations of model sensitivity to humidity in realistic climates and are not intended to be predictive distribution maps.

## Discussion

Here we demonstrate that relative humidity interacts with temperature to alter the performance of key adult traits (e.g., survivorship, fecundity) for an important malaria vector, *Anopheles stephensi*. Although there has been previous work in the literature studying the effects of relative humidity on the thermal sensitivity of these adult traits (e.g., reviewed in Brown et al., 2023) and insects at large (Gol’berg, 1982; Rocha et al., 2001; Yadav and Chaudhary, 1986), this literature has treated humidity as a modifying stressor rather than as a factor that affects thermal performance curves or population-level outcomes. Further, previous work in this field has been limited by a narrow range of environmental conditions, challenging the incorporation of these relationships into mathematical models to quantify thermal performance curves, and spatial and temporal dynamics. Consequently, how relative humidity shapes the thermal performance of adult traits and downstream population fitness in mosquitoes, and insects more broadly, has remained largely unexplored. In this study, we consider the effects of relative humidity on the thermal performance of adult traits and find relative humidity to have varied effects of trait thermal performance. Collectively, these results highlight humidity as a potentially important but underappreciated driver of population dynamics in mosquitoes and insects more broadly.

The main effects of temperature on adult traits were consistent with previous work such that adult lifespan, fecundity, and biting rate exhibited a non-linear, unimodal response to temperature, were constrained by cool (*T*_min_) and warm (*T*_max_) sub-optimal temperatures, and maximized (*T*_pk_) at an intermediate temperature (Brown et al., 2004; Deutsch et al., 2008; Mordecai et al., 2013). Adult traits varied in their shape (symmetric vs. asymmetric), performance breadth (narrow range vs. wide range), as well as the temperatures that optimized performance (Mordecai et al., 2019). In comparing previous work in this system at relative humidity conditions commonly used in laboratory experiments, our TPC predictions for adult traits reflected previous work under certain contexts. For example, Villena et al. (2022), using data aggregated from multiple studies, had distinct predicted thermal peaks for lifespan, daily egg production, and daily biting rate compared to those reported in our study at intermediate relative humidity (60% RH). Instead, our TPC results at close to equivalent relative humidity (75% RH) for adult traits align more closely with Miazgowicz et al. (2020). This is likely because of the use of synthesized data for TPCs (Villena et al., 2022), aggregated data from multiple *Anopheles* species (Mordecai et al., 2013; Johnson et al., 2015), and experimental design considerations such as estimating fecundity and biting phenotypes from the first gonotrophic cycle (Shapiro et al., 2017; Mordecai et al., 2013; Johnson et al., 2015) in these other studies.

Adult lifespan in mosquitoes has long been recognized as a key driver of population dynamics and pathogen transmission, and survival generally declines at warmer temperatures due to accelerated aging and the increased metabolic demands (Barr et al., 2024). Relative humidity further modulates thermal tolerance, particularly in small-bodied ectotherms like mosquitoes, which experience rapid heat and water exchange due to high surface-area-to-volume ratios, cuticular permeability, and spiracular control (Benoit and Denlinger, 2010; Kleynhans and Terblanche, 2011). Empirical and theoretical studies show that water loss rates increase exponentially with temperature and are strongly influenced by atmospheric moisture, such that low humidity accelerates dehydration, elevates metabolic demand, and increases mortality (Gray and Bradley, 2005). Our results are largely consistent with such expectations. Mosquitoes housed in high humidity were found to tolerate warmer temperatures compared to mosquitoes housed in dry conditions.

Despite this rescuing effects of humidity at warmer temperatures, we also observed a decrease in lifespan at optimal temperatures when humidity was high. Similar to temperature, the relationship between organismal fitness and optimal hydroregulation is non-linear, with significant costs to fitness occurring under both dehydrating and overhydrating environments (Anderson and Andrade, 2017; Mitchell and Bergmann, 2016). This could result in complex effects on the thermal performance of insect phenotypes as observed here, with several potential explanations for this effect. First, mosquitoes may rely on low-level evaporative water loss for heat dissipation (Lahondère and Lazzari, 2012). At optimal thermal conditions and high humidity, limited evaporative cooling could accelerate aging through subtle elevations in body temperature and metabolic rate. Second, high humidity promotes microbial growth on cuticle surfaces and in rearing environments. Sublethal infections or increased immune activation could impose chronic energetic costs that reduce lifespan (Ardia et al., 2012). Third, at moderate temperatures, mosquitoes actively regulate spiracular openings to balance oxygen uptake and water conservation. High humidity may relax spiracular control, increasing oxygen consumption and metabolic rate, which can accelerate senescence (Chown, 2002). Finally, we were unable to quantify the effects of relative humidity on lifespan at cooler temperatures nor estimate the *T*_min_ at various humidity levels because we did not collect lifespan data below 16*^◦^*C. More work at lower temperatures or the use of a hierarchical model with a different functional form may improve the uncertainty of our estimates of lifespan and the thermal minima at varying relative humidity.

Previous studies examining humidity effects on mosquito reproduction generally report increased fecundity at higher relative humidity, with low humidity delaying oviposition and reducing egg production (Costa et al., 2010). However, most estimates are based on the first gonotrophic cycle or fecundity estimated from truncated periods of the adult lifespan, which can overestimate lifetime reproductive potential by capturing early-life reproductive peaks (Miazgowicz et al., 2020). Indeed, Miazgowicz et al. (2020) showed that fecundity estimated from early egg production yielded a much higher upper thermal limit than directly observed lifetime fecundity, reflecting rapid reproductive senescence at high temperatures. By quantifying egg production across the entire adult lifespan, our results show that humidity alters both magnitude of daily egg production and its thermal dependence. Dry conditions shifted the thermal optimum for daily egg production toward warmer temperatures but sharply reduced adult lifespan, constraining the duration of reproduction. This pattern is consistent with a life-history trade-off in which females under dry conditions increase short-term reproductive investment at the expense of somatic maintenance, in line with terminal investment theory (Hudson et al., 2020). Such trade-offs are well documented in insects and can be driven by dehydration and heat stress, which accelerate oogenesis and oviposition while elevating metabolic costs, water loss, oxidative stress, and reproductive senescence (Gray and Bradley, 2005; Benoit and Denlinger, 2010; Martin et al., 2024). Consequently, elevated daily fecundity under warm, dry conditions likely reflects faster egg production coupled with a shortened reproductive lifespan, ultimately limiting lifetime fecundity.

We originally hypothesized that relative humidity would influence the thermal performance of the daily biting rate of individual females. Reduced humidity could increase feeding propensity, particularly at warm temperatures, if blood meals serve as an important water source when lifespan is constrained (Hagan et al., 2018a), or alternatively decrease biting rates if mosquitoes reduce activity to limit desiccation, as observed in other insects (Chown and Davis, 2003; Yu et al., 2010). Contrary to these expectations, we detected no clear effect of relative humidity on per capita daily biting rate. This apparent insensitivity likely reflects aspects of our laboratory design rather than a lack of biological responsiveness. We observed substantial among-individual variation in feeding propensity that obscured temperature–biting rate relationships at each humidity level. In addition, females were provided daily access to blood meals, eliminating host-seeking costs and supplying both nutrients and water, which likely buffered dehydration and masked humidity-driven variation in feeding behavior. In natural systems, mosquitoes experience continual water loss and must balance the hydration benefits of blood feeding against the elevated metabolic and water-loss costs of host seeking, particularly under dry conditions (Hagan et al., 2018a). Empirical studies show that both high and low humidity can increase biting activity depending on context. High humidity can enhance sensory responsiveness (Eiras and Jepson, 1994), while dry conditions can increase dehydration-driven host seeking (Khan and Maibach, 1971; Hagan et al., 2018b). However, severe dryness has also been shown to suppress activity to limit water loss (Kessler and Guerin, 2008). The opposing pressures of dehydration and water loss associated with active host-seeking likely generate nonlinear, context-dependent biting responses across temperature-humidity gradients that were not captured under laboratory conditions. What we can conclude is that relative humidity variation is not changing the effects of temperature on the underlying physiological desire to imbibe blood. Additional experiments simulating more realistic host-seeking opportunities in the lab or field with potentially larger sample sizes may provide further resolution on the effects of relative humidity on the daily biting rates in this system.

Integrating the effects of relative humidity on the thermal performance of adult traits into a key metric of population fitness (*r*_m_), demonstrates that relative humidity profoundly altered the thermal performance of *An. stephensi*. Reduced humidity shifted the thermal optimum of *r*_m_ toward warmer temperatures, an effect driven largely by humidity-induced increases in daily fecundity and, to a lesser extent, modest changes in adult lifespan. Sensitivity analyses demonstrated that *r*_m_ is most responsive to variation in fecundity at both high and low temperature extremes, whereas sensitivity to fecundity is comparatively weak at intermediate temperatures. In contrast, the sensitivity of *r*_m_ on adult survival increased steeply at warm temperatures under high humidity but declined sharply under low humidity, consistent with increased desiccation mortality at elevated temperatures. Together, these results indicate that relative humidity modulates the thermal performance of *An. stephensi* population growth rate through opposing mechanisms: humidity both accelerates reproduction and relaxes survival constraints, but dry conditions simultaneously enhance near-term fecundity while truncating reproductive lifespan. Consequently, the net effect of humidity on *r*_m_ reflects the balance between humidity-driven reproductive acceleration and humidity-driven survival costs and suggests that “wetter-is-better” may be overly simplistic when the combined effects of temperature and humidity are considered.

Mapping the sensitivity of *r*_m_ to relative humidity across the native (India) and invasive (Africa) ranges of *An. stephensi* reveals different spatial and seasonal structure in climatic suitability than temperature-only models. While temperature-only projections identify broad regions of high *r*_m_ across tropical Africa and South Asia, incorporating humidity substantially reduces suitability across arid regions (e.g., the Sahara, Sahel, Horn of Africa, and interior Pakistan) and increases *r*_m_ in humid monsoon regions and highland zones. These results extend previous mechanistic models emphasizing temperature as the dominant constraint on mosquito population growth by showing that omission of humidity systematically overestimates both the magnitude and extent of suitable climate space (Miazgowicz et al., 2020; Villena et al., 2022). Seasonal analyses further indicate that humidity enhances *r*_m_ during wet seasons but suppresses it during dry periods, particularly during pre-monsoon conditions in South Asia and African dry seasons, leading to a marked reduction in year-round suitability when humidity is included (Ryan et al., 2023; Acosta et al., 2025). By explicitly linking humidity to adult fitness traits, our trait-based framework provides mechanistic insight into these patterns. Dry conditions favor brief pulses of rapid reproduction but constrain survival, compressing seasonal windows of population growth, whereas moist conditions can partially offset thermal stress under warming. Overall, models incorporating humidity yield distinct estimates of the spatial and temporal dynamics of mosquito population growth and could potentially explain why relative humidity consistently emerges as a strong predictor of malaria transmission (Brown et al., 2023; Acosta et al., 2025).

This study has several limitations. First, experiments were conducted using a long-established *An. stephensi* laboratory colony due to export restrictions from its native range. Laboratory colonies may exhibit founder effects or laboratory adaptation that limit direct inference to locally adapted populations (Ross et al., 2019). However, our goal was not to predict site-specific performance, but to isolate how relative humidity modifies the thermal performance of key mosquito traits. While local adaptation can shift thermal performance parameters (Dennington et al., 2024; Couper et al., 2025), the qualitative form of temperature–trait relationships is generally conserved across mosquito populations and ectotherms (Dennington et al., 2024), and prior comparisons between long-established and field-derived Anopheles colonies show similar thermal responses (Lyons et al., 2012). Nonetheless, extension to natural populations remains an important next step. Second, although our controlled design allowed precise isolation of temperature–humidity interactions, it does not capture environmental complexity such as microclimatic heterogeneity, sub-daily temperature fluctuations, or carryover effects from juvenile environments that may shape adult performance in nature (Murdock et al., 2017; Lambrechts et al., 2011; Paaijmans et al., 2013; Lyons et al., 2013; Shapiro et al., 2017). Finally, habitat availability driven by precipitation, water access, and evaporation will further constrain suitability (Whittaker et al., 2023). Accordingly, our suitability maps should be interpreted as heuristic assessments of how humidity and temperature jointly influence mosquito population fitness, rather than direct predictions of *An. stephensi* distribution or abundance.

In summary, our results show that relative humidity fundamentally reshapes the temperature dependence of key adult fitness traits in *An. stephensi*, altering the thermal performance of lifespan, fecundity, and population growth rate. Rather than simply amplifying temperature effects, humidity shifts thermal optima, modifies performance breadths, and changes the relative importance of traits across environmental gradients. Incorporating these effects into demographic models substantially alters projections of climatic suitability, reducing predicted population growth in hot, arid regions while reshaping the spatial and seasonal timing of favorable conditions elsewhere. These findings challenge temperature-only projections and highlight moisture availability as an essential, but often overlooked, axis of climatic suitability for disease vectors and insects more broadly.

## Acknowledgments

BLJ acknowledges experimental support from Jamie Scanlan, Cole Huh, Nancy-Selwood Metcalfe, Steven Dai, and Ali Mobeen, CCM would like to thank the National Institute of Allergies and Infectious Diseases (NIAID) R01AI163444 and a Climate Health Supplement (NIAID) R01AI153444-03S1 for funding this work.

## Data and code availability

All data and code needed to evaluate the conclusions in the paper are present in the paper, the Supplementary Materials, or the project’s GitHub repository found at: https://anonymous.4open.science/r/An-stephensi-juvenile-traits-34B7

## Statement of authorship

CCM, LRJ, MP, MCW, AM, and PJH conceived the study and designed the experiments. BLJ, JJB, BSL, JS performed the experiments and collected data. MCW and ERB created the maps. PJH, LRJ and BDH performed statistical analyses and modeling. BLJ and PJH wrote the manuscript, and all authors contributed substantially to revisions.

