## Supplemental data for "Humidity shapes the thermal niche of *Anopheles stephensi*, an invasive malaria vector"

### Supplementary Equations

#### *Modelling the fecundity rate and biting rate TPCs*

As noted in the main text, we modeled the fecundity rate and biting rate thermal performance curves (TPCs) for each humidity level by fitting the standard Brière model (SE1; implemented in the `bayesTPC` package in R; Briere et al., 1999; Sorek et al., 2025). Fecundity rate ($b_{\max}$ in Eqn. 1) was calculated as the lifetime number of eggs for each individual divided by its adult lifespan. Biting rate was calculated similarly, as the lifetime number of blood meals for each individual divided by its adult lifespan.

$$1/\alpha(T) = q \cdot T \cdot (T - T_{\min}) \cdot \sqrt{(T_{\max} > T) \cdot (T_{\max} - T)} \cdot (T_{\max} > T) \cdot (T > T_{\min}) \tag{SE1}$$

Here, $T$ is temperature in degrees Celsius, $T_{\min}$ is the low temperature (ºC) at which rates become negative, $T_{\max}$ is the high temperature (ºC) at which rates become negative, and $q$ scale parameter that sets the maximum rate of the curve. The temperatures at which trait performance peaks and its value at its peak ($T_{\text{pk}}$ and $B_{\text{pk}}$, respectively; Table 2, Main text) were estimated numerically from the posterior distributions for each humidity level's TPC and summarized by the posterior medians and Highest Posterior Density (HPD) intervals.

#### *Modelling the adult lifespan TPCs*

We parameterised $z$ in Eqn. 2 (Main text) by substituting TPCs fitted to the adult lifespan data for each humidity level. Adult lifespan was modeled using a temperature-dependent median function with a Weibull likelihood, implemented in the `rjags` package (Plummer, 2025) in R. In JAGS, the Weibull likelihood was implemented using the generalized gamma distribution with the exponent fixed at $a = 1$. Weakly informative priors were specified: normal distributions for $\lambda$ (mean 0, variance 10) and $T_{\text{pk}}$ (mean 22, variance 10), and exponential distributions (rate 0.5) for $c$ and the shape parameter.

We selected this approach because adult lifespan was observed across the full range of experimental temperatures, including survival at the highest temperatures tested, making hard lower and upper thermal limits difficult to estimate directly. Consequently, a model with smooth, unbounded tails provides a more realistic representation of the data than models that impose strict cut-offs. We further selected this temperature-dependent median model with Weibull likelihood over a quadratic alternative because it consistently outperformed the quadratic model based on the Widely Applicable Information Criterion (WAIC; Fig. S1).

For each observation $i = 1, \ldots, N_{\text{obs}}$, the temperature-dependent median and rate parameter are defined as:

$$\text{med}_i = \frac{1}{\exp(\lambda)\left(\text{temp}_i - T_{\text{pk}}\right)^2 + c} \tag{SE2}$$

$$\lambda_i = \frac{\text{med}_i}{[\log(2)]^{1/\text{shape}}} \tag{SE3}$$

The observed trait values are modelled as:

$$\text{trait}_i \sim \text{GenGamma}\left(a = 1,\, \theta = 1/\lambda_i,\, \text{shape}\right) \tag{SE4}$$

where $\text{GenGamma}(a, \theta, \text{shape})$ denotes the generalized gamma distribution with exponent $a$, scale parameter $\theta$, and shape parameter shape. Setting $a = 1$ corresponds to the Weibull distribution.

Peak temperature ($T_{\text{pk}}$) and peak lifespan ($B_{\text{pk}}$) were obtained numerically from posterior samples and summarized using posterior medians and Highest Posterior Density (HPD) intervals.

**Figure S1: Model selection for adult lifespan TPCs across humidity**. Adult lifespan TPCs were fitted using Eqns. SE2–SE4, which were then inverted to give $z$ in Eqn. 1 (Main text; Fig. S2). Boxplot horizontal lines represent medians; lower and upper hinges are the 25th and 75th percentiles. Upper whiskers extend from the hinge to the largest value no further than 1.5 × inter–quartile range (IQR) from the hinge. The lower whisker extends from the hinge to the smallest value at most 1.5 × IQR of the hinge. Points represent individual mosquitoes. Insets show the estimates of $T_{\mathrm{pk}}$ and $B_{\mathrm{pk}}$ with credible intervals for each model. These estimates were obtained numerically from the posterior distributions for each humidity level’s fitted curve and summarized using posterior medians and Highest Posterior Density (HPD) intervals.

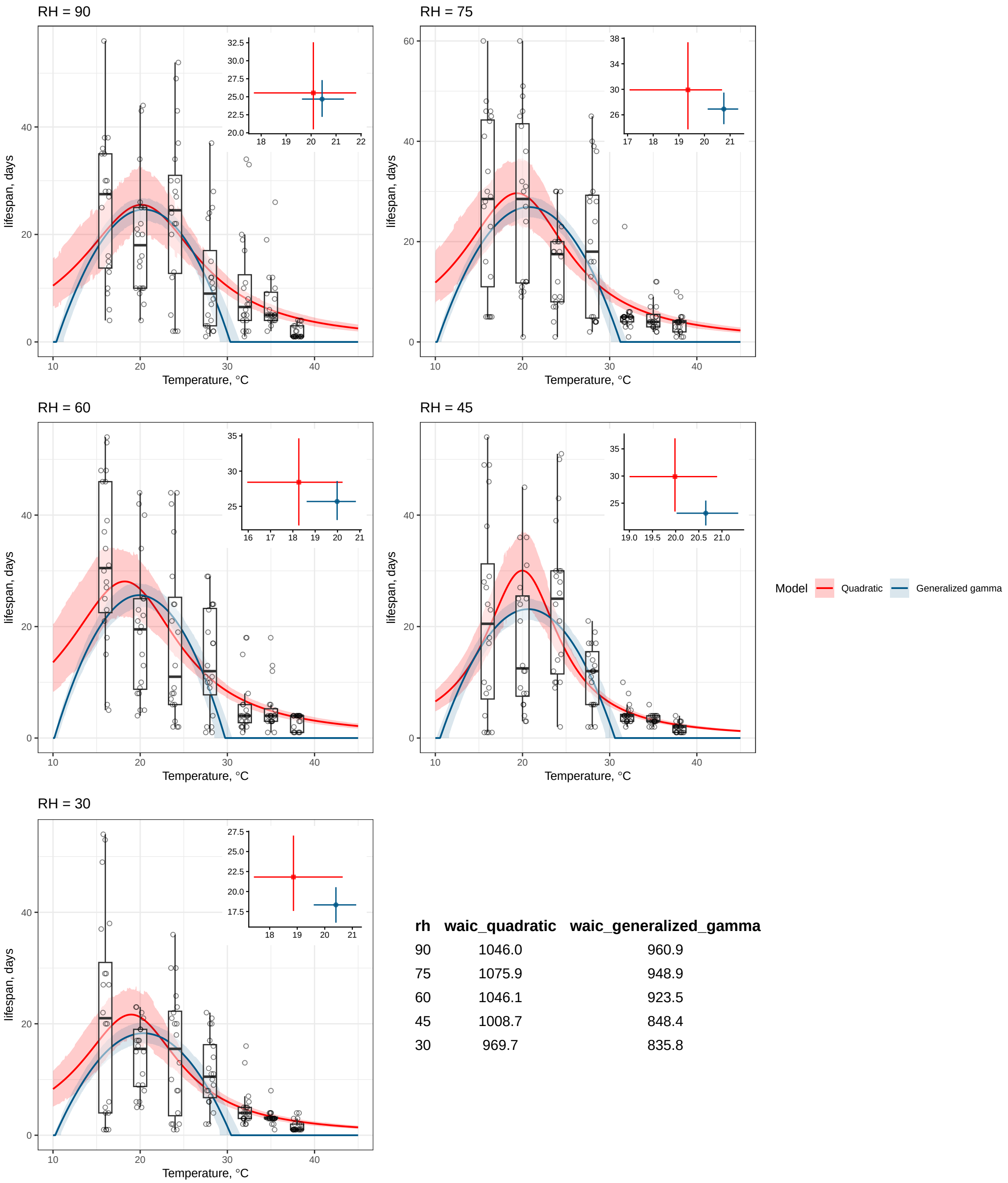


| rh | waic_quadratic | waic_generalized_gamma |
|---|---|---|
| 90 | 1046.0 | 960.9 |
| 75 | 1075.9 | 948.9 |
| 60 | 1046.1 | 923.5 |
| 45 | 1008.7 | 848.4 |
| 30 | 969.7 | 835.8 |

### Trait sensitivity analysis

To determine the extent to which variation in humidity can affect the relative contributions of the individual fitness traits to $r_\mathrm{m}$'s temperature dependence, using the chain rule, we can write (Cator et al., 2020; Mordecai et al., 2013):

$$\frac{\mathrm{d}r_\mathrm{m}}{\mathrm{d}T} = \frac{\partial r_\mathrm{m}}{\partial b_\mathrm{max}}\frac{\mathrm{d}b_\mathrm{max}}{\mathrm{d}T} + \frac{\partial r_\mathrm{m}}{\partial \alpha}\frac{\mathrm{d}\alpha}{\mathrm{d}T} + \frac{\partial r_\mathrm{m}}{\partial z}\frac{\mathrm{d}z}{\mathrm{d}T} + \frac{\partial r_\mathrm{m}}{\partial p_\mathrm{EA}}\frac{\mathrm{d}p_\mathrm{EA}}{\mathrm{d}T} + \frac{\partial r_\mathrm{m}}{\partial \kappa}\frac{\mathrm{d}\kappa}{\mathrm{d}T}. \tag{SE5}$$

Each summed term in the right hand side of this equation quantifies the relative contribution of each trait TPC parameter in Eqn. 2 (Main text) to the temperature dependence of $r_\mathrm{m}$. To calculate the derivatives for each TPC parameter in SE4, we used the MAP (Maximum A Posteriori) estimator for each trait–humidity combination (Figs. 4 & S8). The sample-based MAP estimator is calculated as part of the MCMC fitting process in the `bayesTPC` package (Sorek et al., 2025) in R. We used this method to estimate the Eqn. 2 parameters because it maximizes the unknown parameter's posterior probability, given the observed data and prior beliefs so, compared to the Maximum Likelihood Estimator (MLE), it incorporates prior information.

### Methods for Mapping Study Region and Data Sources

We mapped the maximal mosquito population growth rate ($r_\mathrm{m}$) across South Asia and Africa under historical and future climate conditions. The South Asian study region encompassed India and its neighboring countries (Bangladesh, Bhutan, Nepal, Sri Lanka, Pakistan, Maldives, and Afghanistan), capturing diverse climates from tropical to temperate zones. The African study region covered the entire continent, incorporating equatorial, savannah, arid, and subtropical climates to enable broad regional comparisons of potential mosquito growth.

Climate variables, including daily maximum temperature, minimum temperature, and specific humidity, were obtained from the NASA NEX–GDDP–CMIP6 dataset. These statistically downscaled, bias–corrected projections have a resolution of $0.25^\circ \times 0.25^\circ$ ($\sim$27.8 km). We incorporated outputs from 23 general circulation models (GCMs) in this dataset, analyzing the ensemble of historical (1970–2000) model runs to generate climate inputs for the models.

#### *Climate Data Processing*

We used the NASA climate data to estimate ($r_\mathrm{m}$), which integrates the effects of temperature and humidity on mosquito population dynamics. Daily climate variables including maximum temperature, minimum temperature, and specific humidity from 1970 to 2000 were masked to terrestrial grid cells within the study–region boundaries. These daily data were first aggregated into monthly averages. Monthly mean temperatures were calculated as the average of daily maximum and minimum temperatures. Relative humidity was computed from monthly averaged specific humidity data using the Magnus–Tetens equation, accounting for temperature-dependent saturation vapor pressure. Finally, seasonal means were calculated by averaging monthly data across defined three–month periods (e.g., January—March) for the entire 1970–2000 period.

#### *Trait-Based Population Growth Model*

Monthly mosquito population growth was predicted using the trait-based model of the intrinsic rate of population increase ($r_\mathrm{m}$). Biological traits, including juvenile survival, developmental rate, and fecundity, were modeled as nonlinear functions of temperature based on laboratory experiments. Temperature-trait curves were derived the experimental RH values of 30%, 45%, 60%, 75%, and 90%. To quantify the influence of humidity, we conducted two model runs. The temperature-only model held humidity constant at a fixed reference value of 75% RH, reflecting the baseline condition under which most temperature-trait models are derived. The temperature $\times$ RH model used the actual RH values from the climate dataset. We applied piecewise linear interpolation to handle continuous RH variations in the climate data, deriving each modeled trait from the two nearest experimental RH levels. This method ensured smooth trait responses to varying humidity without abrupt category transitions. These analyses produced monthly raster datasets of $r_\mathrm{m}$ values for the two model runs, and the monthly outputs were summarized to generate mean annual $r_\mathrm{m}$.

#### *Spatial Analysis and Mapping*

Maps of mean annual $r_\mathrm{m}$ were generated to highlight spatial variability in mosquito population growth potential. Difference maps ($\Delta r_\mathrm{m}$) were created to highlight areas where adding humidity altered the growth rates, revealing regions of higher or lower suitability for population growth. We also computed the total land area for which climate was suitable for mosquito growth ($r_\mathrm{m} > 0$) during all months of the year (Fig. S9).

### Supplementary figures and tables

**Table S1:** Estimates for the adult lifespan TPC parameters; $B_{pk}$ and $T_{pk}$. Numerical posterior estimates (median ± 95% credible intervals) of the parameters from $n$ = 10,000 iterations for a burn–in of $n$ = 5000.

| RH (%) | $B_{pk}$ | $B_{pk}$ lower | $B_{pk}$ upper | $T_{pk}$ | $T_{pk}$ lower | $T_{pk}$ upper |
|---|---|---|---|---|---|---|
| 90 | 25.757 | 20.350 | 32.188 | 20.213 | 17.918 | 22.087 |
| 75 | 30.025 | 24.430 | 36.782 | 19.424 | 17.357 | 20.896 |
| 60 | 28.496 | 22.079 | 35.426 | 18.198 | 15.956 | 20.160 |
| 45 | 29.666 | 22.980 | 38.116 | 20.055 | 19.004 | 20.861 |
| 30 | 21.820 | 16.675 | 27.482 | 18.969 | 17.182 | 20.511 |

**Table S2: Estimates for the adult fecundity rate TPC parameters; $T_{min}$ and $T_{max}$.** Numerical posterior estimates (median ± 95% credible intervals) of the parameters from $n$ = 10,000 iterations for a burn–in of $n$ = 5000.

| RH (%) | $T_{min}$ | $T_{min}$ lower | $T_{min}$ upper | $T_{max}$ | $T_{max}$ lower | $T_{max}$ upper |
|---|---|---|---|---|---|---|
| 90 | 14.776 | 12.604 | 16.587 | 36.472 | 35.877 | 37.230 |
| 75 | 10.761 | 10.000 | 12.286 | 38.960 | 38.427 | 39.671 |
| 60 | 10.855 | 10.000 | 12.548 | 38.438 | 38.045 | 38.999 |
| 45 | 13.652 | 10.980 | 16.006 | 39.656 | 38.942 | 40.000 |
| 30 | 18.300 | 15.999 | 19.998 | 38.154 | 37.559 | 39.380 |

**Table S3: Estimates for the adult fecundity rate TPC parameters; $B_{pk}$ and $T_{pk}$.** Numerical posterior estimates (median ± 95% credible intervals) of the parameters from $n$ = 10,000 iterations for a burn–in of $n$ = 5000.

| RH (%) | $B_{pk}$ | $B_{pk}$ lower | $B_{pk}$ upper | $T_{pk}$ | $T_{pk}$ lower | $T_{pk}$ upper |
|---|---|---|---|---|---|---|
| 90 | 30.290 | 28.122 | 32.659 | 31.131 | 30.681 | 31.582 |
| 75 | 29.376 | 27.520 | 31.171 | 32.482 | 31.982 | 32.983 |
| 60 | 26.026 | 24.185 | 28.069 | 32.082 | 31.732 | 32.533 |
| 45 | 32.081 | 29.687 | 34.556 | 33.383 | 32.733 | 33.884 |
| 30 | 28.185 | 25.358 | 31.016 | 33.083 | 32.432 | 33.884 |

**Table S4: Estimates for the biting rate TPC parameters; $T_{min}$ and $T_{max}$.** Numerical posterior estimates (median ± 95% credible intervals) from $n$ = 10,000 iterations with a burn–in of $n$ = 5000.

| RH (%) | $T_{min}$ | $T_{min}$ lower | $T_{min}$ upper | $T_{max}$ | $T_{max}$ lower | $T_{max}$ upper |
|---|---|---|---|---|---|---|
| 90 | 12.957 | 10.001 | 16.944 | 39.233 | 37.839 | 39.999 |
| 75 | 14.503 | 10.417 | 18.497 | 39.640 | 38.900 | 40.000 |
| 60 | 11.175 | 10.001 | 14.029 | 39.101 | 37.588 | 40.000 |
| 45 | 13.719 | 10.308 | 16.764 | 39.450 | 38.521 | 40.000 |
| 30 | 11.537 | 10.000 | 15.095 | 38.801 | 35.899 | 40.000 |

**Table S5: Estimates for the biting rate TPC parameters; $B_{pk}$ and $T_{pk}$.** Numerical posterior estimates (median ± 95% credible intervals) from $n$ = 10,000 iterations with a burn–in of $n$ = 5000.

| RH (%) | $B_{pk}$ | $B_{pk}$ lower | $B_{pk}$ upper | $T_{pk}$ | $T_{pk}$ lower | $T_{pk}$ upper |
|---|---|---|---|---|---|---|
| 90 | 0.324 | 0.280 | 0.374 | 32.983 | 31.732 | 33.884 |
| 75 | 0.316 | 0.265 | 0.365 | 33.483 | 32.733 | 34.234 |
| 60 | 0.307 | 0.268 | 0.344 | 32.683 | 31.431 | 33.584 |
| 45 | 0.341 | 0.299 | 0.388 | 33.233 | 32.432 | 33.984 |
| 30 | 0.298 | 0.246 | 0.343 | 32.482 | 30.130 | 33.584 |

**Table S6: Estimates for the $r_m$ TPC parameters; $T_{opt}$ and $r_{opt}$.** Numerical posterior estimates (median ± 95% credible intervals) of the $r_m$ model (Eqn. 1, Main text) parameters from $n$ = 10,000 iterations for a burn–in of $n$ = 5000.

| RH (%) | $r_{opt}$ | $r_{opt}$ lower | $r_{opt}$ upper | $T_{opt}$ | $T_{opt}$ lower | $T_{opt}$ upper |
|---|---|---|---|---|---|---|
| 90 | 0.3597 | 0.3431 | 0.3756 | 32.7828 | 32.5325 | 33.0831 |
| 75 | 0.3950 | 0.3795 | 0.4091 | 34.8348 | 34.5846 | 35.1351 |
| 60 | 0.3809 | 0.3688 | 0.3938 | 34.2342 | 33.9339 | 34.4845 |
| 45 | 0.4068 | 0.3964 | 0.4171 | 34.8849 | 34.4845 | 35.1351 |
| 30 | 0.3732 | 0.3621 | 0.3845 | 35.1351 | 34.6346 | 35.6356 |

**Table S7: Estimates for the $r_m$ TPC parameters; $T_{min}$ and $T_{max}$.** Numerical posterior estimates (median ± 95% credible intervals) of the parameters from $n$ = 10,000 iterations for a burn–in of $n$ = 5000.

| RH (%) | $T_{min}$ | $T_{min}$ lower | $T_{min}$ upper | $T_{max}$ | $T_{max}$ lower | $T_{max}$ upper |
|---|---|---|---|---|---|---|
| 90 | 15.1652 | 13.8138 | 16.5666 | 36.4364 | 35.8358 | 37.1371 |
| 75 | 14.6146 | 14.1642 | 15.0150 | 38.9389 | 38.3884 | 39.6396 |
| 60 | 15.0150 | 14.5646 | 15.4655 | 38.3884 | 38.0380 | 38.9389 |
| 45 | 15.0651 | 14.1141 | 16.1662 | 39.6396 | 38.8889 | 39.9399 |
| 30 | 18.7187 | 17.3173 | 20.0701 | 38.1381 | 37.5375 | 39.4895 |

**Figure S2: Adult mortality rate TPCs (1/adult lifespan) used for the temperature– and humidity–dependent $r_m$ calculations.** Adult lifespan TPCs were fitted using Eqns. SE2–SE4 which were then inverted to give $z$ in Eqn. 1 (Main text). Relative humidity (%) levels are shown in the legend.

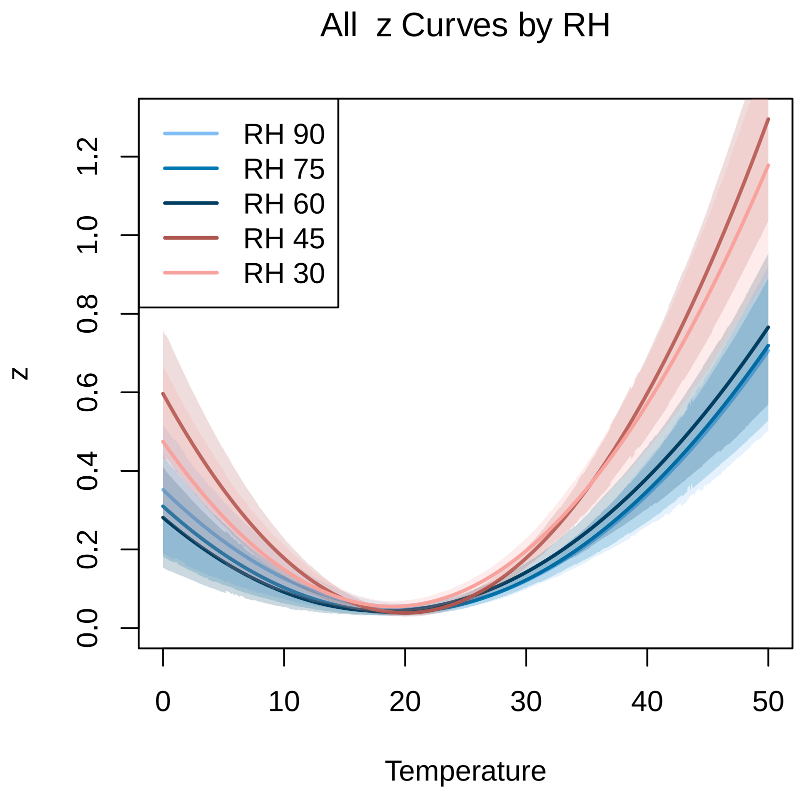


Adult lifespan TPCs were fitted using Eqns. SE2–SE4
